# CellOntologyMapper: Consensus mapping of cell type annotation

**DOI:** 10.1101/2025.06.10.658951

**Authors:** Zehua Zeng, Xuehai Wang, Hongwu Du

## Abstract

Single-cell RNA sequencing has revolutionized cellular biology, with atlases now encompassing over 100 million cells. However, researchers employ vastly different naming conventions when annotating cell types, creating a fragmented landscape that severely impedes data integration and comparative analysis. Here, we present CellOntologyMapper, an automated framework that standardizes cell type annotations by intelligently mapping user-defined names to established Cell Ontology and Cell Taxonomy identifiers. Our approach leverages advanced natural language processing, including sentence transformers and large language models, to interpret diverse naming conventions and resolve them to standardized ontological terms. The system handles complex challenges including abbreviated cell names, synonym resolution, and context-dependent interpretation. We built a comprehensive query system based on 19,381 cell type entries from established Cell Ontology and Cell Taxonomy databases, which organize into 24 biologically coherent clusters. Systematic validation across datasets spanning different scales— from 17-cell-type lung studies to 91-cell-type immune atlases and complex developmental systems—consistently demonstrated robust performance and high accuracy. CellOntologyMapper successfully resolved annotation challenges across conventional tissue studies, rare cell populations, and developmentally intricate datasets rich in abbreviated nomenclature. By providing automated, scalable annotation harmonization, our framework enables researchers to leverage existing single-cell datasets while ensuring compatibility with future atlas efforts. When incorporated into publications, CellOntologyMapper enhances the credibility and reference value of cell type annotations, representing a crucial step toward an integrated single-cell genomics ecosystem.

## Introduction

Single-cell RNA sequencing (scRNA-seq) has revolutionized our understanding of cellular diversity across tissues, developmental stages, and disease states. As single-cell atlases continue to expand—with some now encompassing over 100 million cells—accurate cell type annotation has emerged as a fundamental prerequisite for meaningful biological interpretation and cross-study integration^[1, 2]^.

Despite this critical importance, the single-cell genomics field faces a persistent challenge: researchers employ vastly different naming conventions when annotating cell types, creating a fragmented landscape that impedes data integration and comparative analysis^[3]^. While individual studies may use descriptive names like “inflammatory macrophages,” “CD14+ monocytes,” or simply “Mac1,” these labels often refer to overlapping or identical cell populations, making cross-dataset comparisons extraordinarily difficult^[4]^. This challenge becomes even more complex in cross-species studies, where equivalent cell types may be annotated with species-specific nomenclatures that obscure evolutionary relationships and functional conservation patterns^[5]^. Current approaches to address this challenge—re-annotating integrated datasets or manual harmonization—introduce substantial limitations^[6]^. Re-annotation after data integration risks batch effect confounding, while manual harmonization is labor-intensive, subjective, and scales poorly with expanding data volumes^[7]^.

The scientific community has developed standardized ontologies, notably the Cell Ontology (CL) and Cell Taxonomy databases, which provide hierarchical, controlled vocabularies for cell type classification^[8]^. However, adoption remains inconsistent across the research community, with many studies continuing to use laboratory-specific nomenclatures^[9]^.

To bridge this critical gap, we developed CellOntologyMapper (https://github.com/Starlitnightly/CellOntologyMapper), a computational framework that automatically standardizes cell type annotations by mapping user-defined cell names to established Cell Ontology^[10]^ and Cell Taxonomy^[11]^ identifiers. Our approach leverages state-of-the-art natural language processing techniques, including advanced sentence transformers and large language models, to intelligently interpret diverse naming conventions and resolve them to standardized ontological terms. The system handles common challenges including abbreviated cell names, synonym resolution, and context-dependent interpretation based on tissue origin and experimental conditions^[8]^. The framework is implemented as an accessible Python package within the OmicVerse^[12]^ ecosystem, designed to integrate seamlessly into existing single-cell analysis workflows.

## Result and Discussion

### Overview of CellOntologyMapper design

To address the challenge of inconsistent cell type naming across single-cell studies, we developed CellOntologyMapper, a comprehensive framework that standardizes cell type nomenclature through two interconnected modules (Figure 1a-b). The first module constructs a robust Cell Ontology Query Database by extracting cell type names from authoritative Cell Ontology and Cell Taxonomy databases along with their associated metadata^[13]^. We employed state-of-the-art sentence transformer models, including FlagEmbedding (BAAI) and Qwen (Alibaba), to generate high-dimensional embedding vectors that capture semantic relationships between cell type names^[14, 15]^. These embeddings, coupled with their corresponding cell type identifiers, are efficiently stored in an SQL database optimized for rapid similarity searches (Figure 1a).

**Figure 1.**
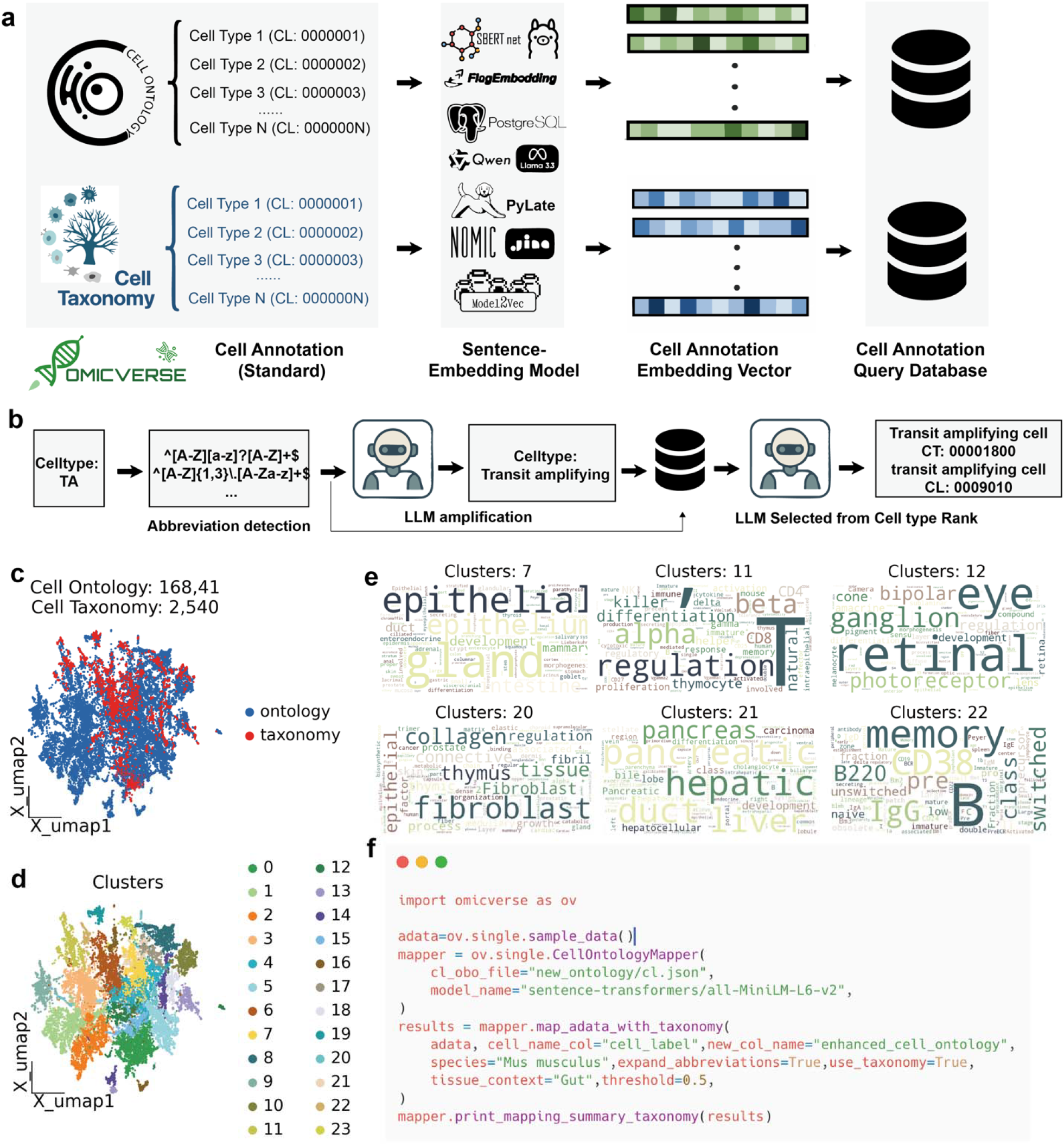
Overview of CellOntologyMapper for Comprehensive Cell Type Standardization. (a) Workflow of constructing the Cell Ontology Query Database. Cell type names were extracted from Cell Ontology and Cell Taxonomy databases, embedding vectors were generated using state-of-the-art sentence transformers (FlagEmbedding, Qwen), and subsequently stored in an SQL database for efficient querying. (b) Pipeline for mapping user-provided cell type abbreviations (e.g., “TA”) to standardized names using regular expression (regex) detection followed by Large Language Model (LLM) amplification. The LLM suggests full cell type names based on sequencing context and tissue background, computes their embedding vectors, ranks similarities against database entries, and selects appropriate Cell Ontology (CL) and Cell Taxonomy (CT) identifiers from top candidates. (c) Summary of the constructed query database containing 19,381 cell type entries (16,841 from Cell Ontology, 2,540 from Cell Taxonomy), visualized on a UMAP embedding to illustrate the distribution of database entries. (d) Unsupervised clustering of cell type embeddings from the query database. Principal component analysis (PCA), neighborhood graph construction, and Leiden clustering identified 24 distinct clusters, each color-coded. (e) Word cloud visualizations highlighting representative cell types from selected clusters. Cluster 7 primarily includes epithelial-related cell types; Cluster 11 is enriched in T cells and subtypes; Cluster 12 consists predominantly of retinal-related cells; Cluster 20 represents fibroblast-related entries; Cluster 21 groups cells associated with liver and pancreas; and Cluster 22 predominantly includes B cells and their subtypes. (f) Example code snippet demonstrating the straightforward and user-friendly API provided by OmicVerse for efficient cell type annotation mapping.

The second module implements an intelligent mapping pipeline that transforms user-provided cell type names into standardized ontology terms. To handle the prevalent issue of abbreviated cell type names, we first apply regular expression-based detection to identify potential abbreviations. When abbreviations are detected, we leverage large language models (LLMs) to generate contextually appropriate full cell type names based on experimental metadata such as tissue origin and sequencing protocol^[16, 17]^. The system then computes embedding vectors for the query terms and performs similarity ranking against the entire database. Finally, an LLM-assisted selection process identifies the most appropriate Cell Ontology and Cell Taxonomy identifiers from the top-ranked candidates, ensuring both semantic accuracy and biological relevance (Figure 1b).

Our comprehensive query database encompasses 19,381 curated cell type entries, comprising 16,841 from Cell Ontology and 2,540 from Cell Taxonomy (Figure 1c). To investigate the semantic organization of this database, we performed dimensionality reduction using UMAP and applied unsupervised Leiden clustering, revealing 24 distinct clusters that group semantically related cell types (Figure 1d). Remarkably, these clusters exhibit clear biological coherence: cluster 7 predominantly contains epithelial cell types, cluster 11 is enriched for T cells and their specialized subtypes, cluster 12 groups retinal cell populations, cluster 20 encompasses fibroblast-related entries, cluster 21 aggregates hepatic and pancreatic cell types, and cluster 22 primarily consists of B cell lineages and their developmental stages (Figure 1e). This clustering pattern validates the biological meaningfulness of our embedding approach and demonstrates the system’s capacity to capture nuanced relationships within the cellular taxonomy. To ensure broad accessibility, we provide a streamlined, user-friendly API through the OmicVerse package that enables researchers to perform cell type standardization with minimal code requirements (Figure 1f). This implementation facilitates seamless integration into existing single-cell analysis workflows while maintaining the sophisticated mapping capabilities of the underlying framework.

### CellOntologyMapper demonstrates robust cross-scale performance across diverse tissue contexts and annotation complexities

To rigorously evaluate the versatility and precision of CellOntologyMapper, we conducted comprehensive validation across datasets spanning dramatically different scales and biological contexts, from focused tissue-specific studies to complex developmental systems rich in specialized nomenclature. This systematic assessment encompassed small-scale datasets (<20 cell types)^[18]^, medium-scale studies (20-50 cell types), large-scale atlases (>50 cell types), and challenging datasets characterized by extensive abbreviations or intricate developmental terminology^[19, 20]^. We first tested CellOntologyMapper on a focused lung tissue dataset (Vieira Braga et al.) containing 17 well-characterized cell types, including immune populations, structural cells, and specialized pulmonary cell lineages^[21]^. The system achieved complete accuracy in mapping all annotated cell types to their corresponding Cell Taxonomy identifiers, successfully distinguishing between closely related populations such as alveolar macrophages, interstitial macrophages, and dendritic cell subsets (Figure 2a). This performance demonstrates the tool’s precision in handling conventional cell type annotations within established tissue contexts.

**Figure 2.**
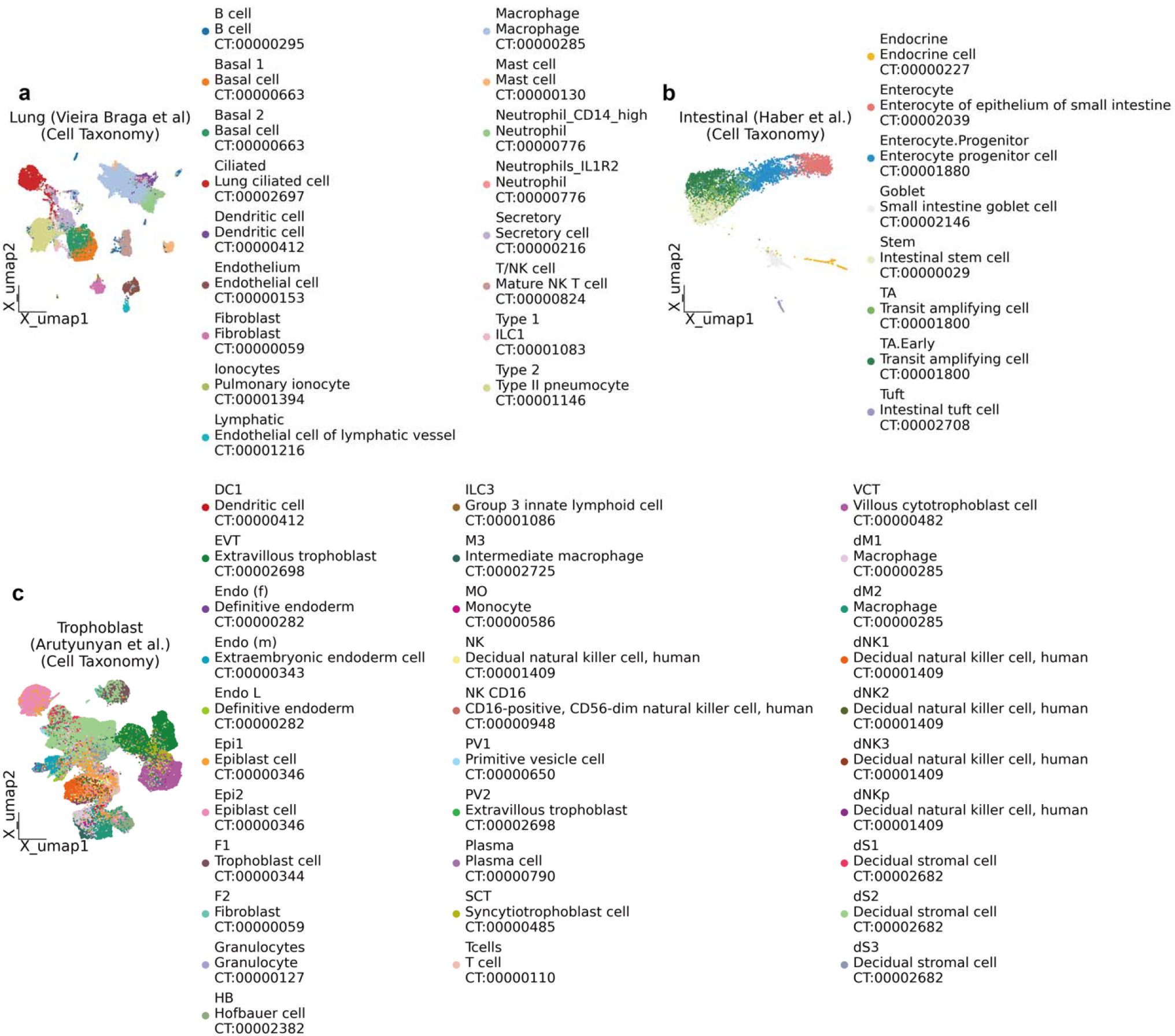
Cell Taxonomy-based Annotation of Single-cell Datasets Across Multiple Tissues. (a) UMAP visualization of annotated lung single-cell dataset (Vieira Braga et al.) based on Cell Taxonomy. Identified cell types include immune cells (e.g., B cells, macrophages, mast cells), structural cells (basal cells, fibroblasts), and specialized epithelial cells (ciliated, secretory). (b) Annotation of intestinal single-cell data (Haber et al.) visualized by UMAP, highlighting differentiation stages and specialized epithelial populations such as enterocytes, transit amplifying cells (TA), goblet cells, and endocrine cells. (c) UMAP-based annotation of trophoblast single-cell dataset (Arutyunyan et al.) demonstrating diverse placental cell populations including extravillous trophoblasts (EVT), decidual stromal cells (dS), natural killer cells (NK), and syncytiotrophoblasts (SCT), annotated comprehensively via Cell Taxonomy (CT) identifiers.

To challenge the system with greater biological complexity, we applied CellOntologyMapper to an intestinal epithelial dataset (Haber et al.) ^[18]^ characterized by extensive cellular differentiation gradients and rare specialized populations. This dataset presented unique annotation challenges, including multiple intermediate differentiation states, region-specific epithelial variants, and highly specialized cell types such as tuft cells and enteroendocrine subtypes. CellOntologyMapper successfully navigated this complexity, accurately annotating diverse populations from proliferative transit-amplifying cells to terminally differentiated absorptive enterocytes, while correctly identifying rare populations like Paneth cells and goblet cell subtypes (Figure 2b). This performance highlights the system’s capacity to handle nuanced biological contexts where cell identity exists along continuous differentiation spectra.

The most stringent test involved a developmentally complex trophoblast dataset (Arutyunyan et al.)^[22]^, which combines the challenges of embryonic cell nomenclature with extensive use of abbreviated cell type names. This dataset encompasses intricate placental development lineages, including syncytiotrophoblasts, cytotrophoblasts, and various extravillous trophoblast populations, many of which are commonly referenced by acronyms in the literature. CellOntologyMapper demonstrated exceptional performance in this challenging context, successfully resolving ambiguous abbreviations and mapping each population to validated Cell Taxonomy entries while maintaining biological coherence across the developmental trajectory (Figure 2c). The system’s ability to distinguish between closely related developmental stages, such as different trophoblast progenitor states, underscores its sophisticated understanding of both semantic relationships and biological context.

Extended validation across additional datasets of varying scales—including medium-scale bone marrow studies (24 cell types), large-scale bone atlases (55 cell types), and comprehensive immune cell compendiums (91 cell types)—consistently demonstrated high accuracy and computational scalability. These results collectively establish CellOntologyMapper as a robust solution capable of handling the full spectrum of single-cell annotation challenges, from straightforward tissue-specific studies to the most complex developmental and immune system atlases encountered in contemporary single-cell genomics.

*Supplementary Figure S1* | *Cell-Taxonomy-based annotation of human bone-marrow single-cell transcriptomes (Zeng et al*.)

*(a) UMAP projection of 34,315 bone-marrow cells annotated with Cell Taxonomy (CT) identifiers at a coarse resolution. Distinct clusters correspond to early progenitors (HSC, MPP, CLP, GMP, CMP), lineage-restricted progenitors (MEP, Eo/Baso/Mast precursor, GMP-Neut, GMP-Mono) and mature immune populations including B cells, naïve and memory CD4L/CD8L T cells, NK cells, monocytes, cDCs, pDCs and plasma cells. A small stromal fraction is tentatively labelled “Stromal cell of bone marrow”, yet the transcriptional separation from erythro-myeloid progenitors is modest, cautioning against over-interpretation of this assignment*.

*(b) The identical dataset re-annotated at higher resolution illustrates the hierarchical depth available in the taxonomy. Additional T-cell states emerge—central-memory, effector-memory, tissue-resident and activated subsets—together with discrete stages of B-cell development (immature B, large pre-B, early and late pro-B). Granular erythro-megakaryocytic differentiation (pro-erythroblast, orthochromatic erythroblast, megakaryocyte) and proliferative intermediates (“Cycling progenitor”, “Proliferative T”) are also resolved. Notably, several clusters assigned as “Pre-B” versus “Pro-B cycling” have overlapping marker repertoires, implying that the apparent separation may partly reflect cell-cycle effects rather than true developmental bifurcation*.

*Across both panels, coloured bullets denote CT IDs (CT0000xxxx), providing an explicit, machine-readable linkage to the underlying ontology. While the taxonomy offers a reproducible framework, users should scrutinise clusters with marginal transcriptional distances—especially where cell-cycle or technical variation could masquerade as biological identity*.

*Supplementary Figure S2* | *Cell-Taxonomy-based annotation of the cross-tissue human immune-cell atlas (Domínguez Conde et al*.)

*(a) Uniform Manifold Approximation and Projection (UMAP) of 79 k single cells profiled across 16 tissues and re-labelled with Cell Taxonomy (CT) identifiers. Sixty transcriptionally distinct clusters are recovered. Lymphoid lineages comprise (i) progressive B-cell maturation from immature-B and transitional-B through germinal-centre (GC) B and plasma cells; (ii) a continuum of T-cell states—naïve, central-memory (TCM), effector-memory (TEM), Temra, tissue-resident, follicular-helper (Tfh), regulatory (Treg), γδ-T, mucosal-associated invariant T (MAIT) and NKT cells; and (iii) three innate-lymphoid-cell branches (ILC1–3) in addition to CD56^bright and CD56^dim NK subsets. The myeloid compartment resolves classical, intermediate and non-classical monocytes, neutrophil–myeloid progenitors, conventional dendritic-cell axes (cDC1, cDC2, DC3), LAMP3^+ migratory DCs, macrophage specialisations (Kupffer, kidney-resident, alveolar) and mast cells. Haematopoietic stem and multipotent progenitors (HSC/MPP), lineage-restricted erythro-megakaryocytic (MEP) and granulocyte-monocyte (GMP) precursors, together with sporadic stromal contaminants—endothelial, epithelial and fibroblast signatures—are also detected*.

## Conclusion

CellOntologyMapper addresses provide the first comprehensive, automated framework for standardizing cell type nomenclature through intelligent mapping to established ontological databases. Our systematic validation across diverse biological contexts demonstrates robust performance regardless of dataset scale or annotation complexity. The significance of CellOntologyMapper extends beyond technical convenience to enable transformative advances in single-cell biology. By resolving annotation fragmentation, our framework unlocks the potential for large-scale meta-analyses. This capability is particularly crucial as the field constructs comprehensive cellular atlases spanning multiple species, developmental stages, and disease conditions. The tool’s sophisticated handling of abbreviated nomenclatures and biological context ensures compatibility with diverse annotation practices across research communities.

Our implementation within the OmicVerse ecosystem provides accessible, user-friendly annotation standardization that will accelerate adoption across the single-cell community. When researchers incorporate CellOntologyMapper results into their publications, their cell type annotations gain enhanced credibility, broader applicability, and improved reference value for the scientific community. As single-cell atlases continue expanding and the volume of cellular data grows exponentially, automated annotation standardization will become essential infrastructure for maintaining scientific rigor and enabling meaningful cross-study comparisons. CellOntologyMapper represents an important step toward a truly integrated single-cell genomics ecosystem that maximizes the scientific value of our collective cellular discoveries.

## Author Contributions

**Zehua Zeng**: Conceptualization; methodology; software; data curation; investigation; validation; writing—original draft; formal analysis; visualization. **Xuehai Wang**: methodology; software; data curation; investigation; **Hongwu Du**: Conceptualization; Writing—review and editing.

