## supplementary for "CellOntologyMapper: Consensus mapping of cell type annotation"

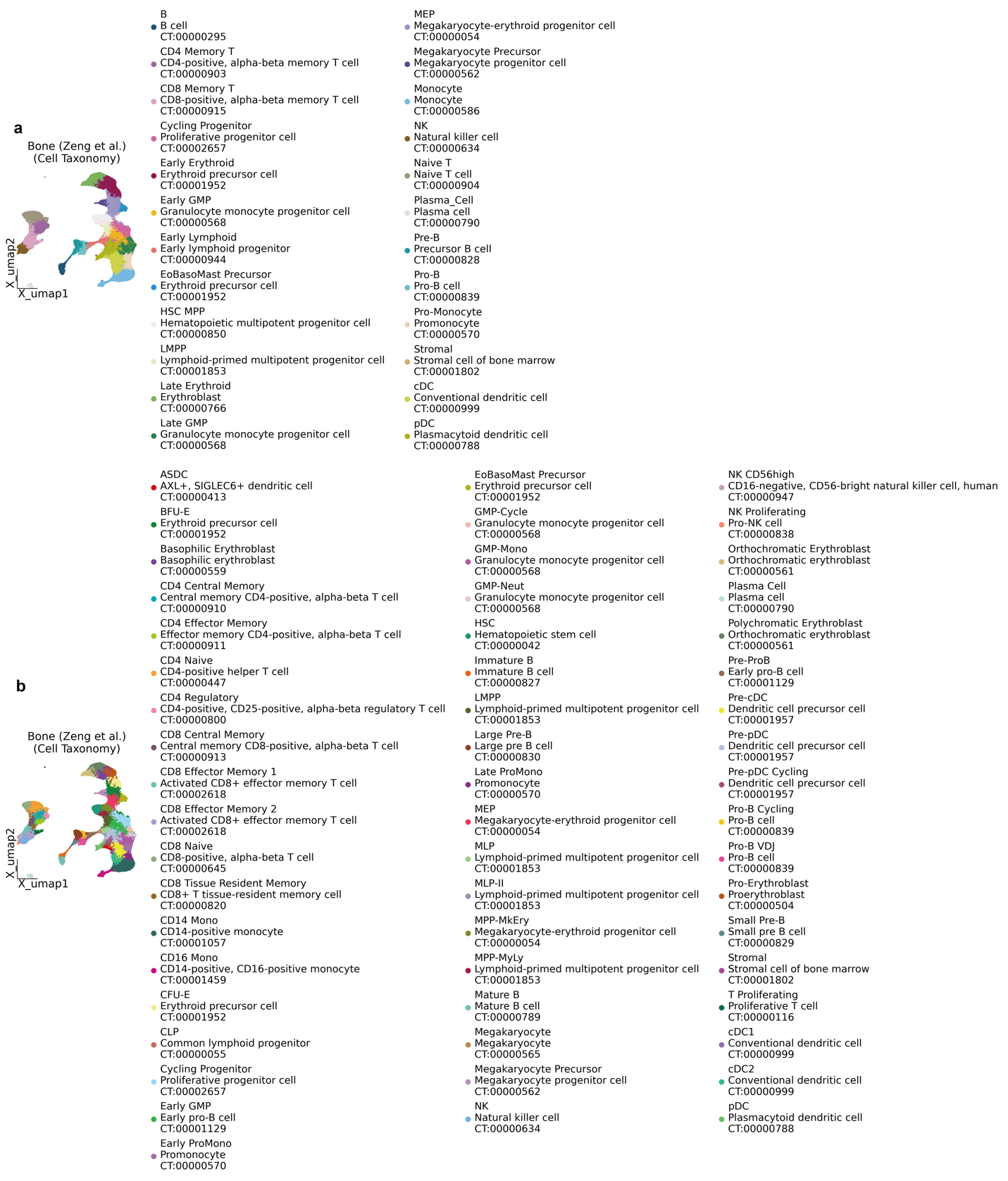

sFigure 1 | Cell Taxonomy-based Annotation of Single-cell Datasets Across Median datasets

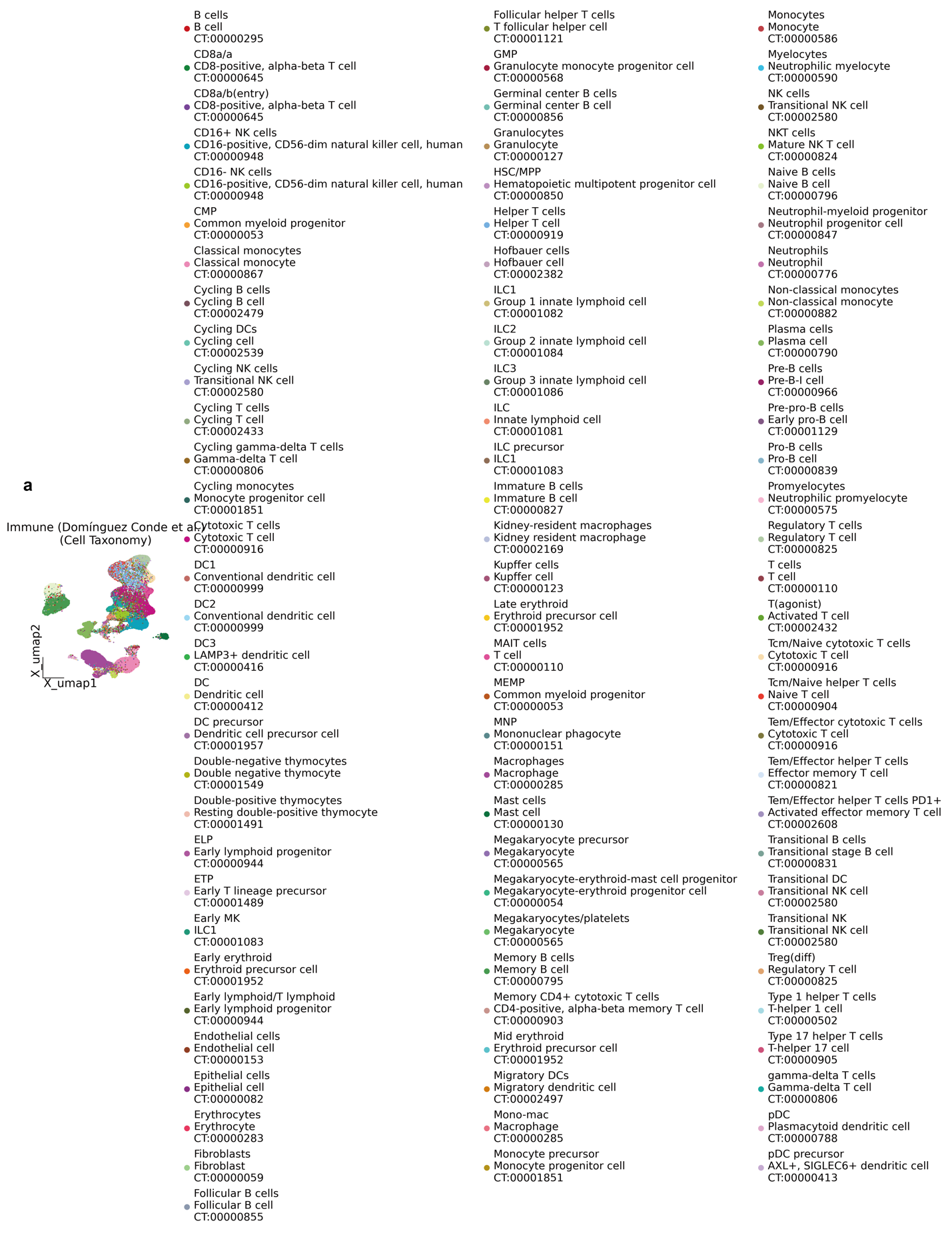

sFigure 2 | Cell Taxonomy-based Annotation of Single-cell Datasets Across Large datasets
